# Metastatic profiling of HER2-positive breast cancer cell lines in xenograft models

**DOI:** 10.1101/2021.10.20.464943

**Authors:** Yuxuan Han, Kazushi Azuma, Shinya Watanabe, Kentaro Semba, Jun Nakayama

## Abstract

Most studies on breast cancer metastasis have been performed using triple-negative breast cancer (TNBC) cells; thus, subtype-dependent metastatic ability of breast cancer is poorly understood. In this research, we performed intravenous injection (IVI) and intra-caudal arterial injections (CAI) using nine human epidermal growth factor receptor-2 (HER2)-positive breast cancer cell lines for evaluating their metastatic abilities. Our results showed that MDA-MB-453, UACC-893, and HCC-202 had strong bone metastatic abilities, whereas HCC-2218 and HCC-1419 did not show bone metastasis. HER2-positive cell lines could hardly metastasize to the lung through IVI. From the genomic analysis, gene signatures were extracted according to the breast cancer subtypes and their metastatic preferences. The UACC-893 cell line was identified as a useful model for the metastasis study of HER2-positive breast cancer. Combined with our previous result on brain proliferation ability, we provide a characteristic metastasis profile of HER2-positive breast cancer cell lines in this study.

**Statements and Declarations:** *Funding:* This study was supported by JSPS KAKENHI (grant no. 18K16269: Grant-in-Aid for Early-Career Scientist to J.N.; grant no. 20J01794, Grant-in-Aid for JSPS fellows to J.N.; grant no. 20J23297, Grant-in-Aid for JSPS fellows to Y.H.) and partially supported by the grants for translational research programs from Fukushima Prefecture (S.W. and K.S.).

*Authorship:* YH and KA performed the *in vivo* experiments and bioinformatical analyses. SW, and KS interpreted the data. YH, KA, and JN wrote the manuscript. JN conceived and designed the study. All the authors reviewed and edited the manuscript.

*Competing Interests:* The authors declare that they have no competing interests.

*Ethical approval:* The animal experiments were conducted under the approval of the ethics committee of Waseda University (2020-A067, 2021-A074).

## Introduction

Breast cancer is the most frequently diagnosed cancer worldwide and appears as the leading cause of cancer death in females [1]. Cancer cell lines derived from human tumors are widely used in metastasis study for their potential usefulness in evaluating preclinical trials [2]. Triple-negative subtype MDA-MB-231 cells and luminal-A subtype MCF7 cells have been most frequently studied in breast cancer metastasis. A recent study that used intracardiac transplantation released a large-scale metastasis map (MetMap500) of human cancer cell lines [3]. The work provided a large-scale characterization of human cancer cell lines and the tool to examine their molecular mechanisms in organ-specific microenvironments. However, their metastasis map was mainly constructed using the groups of luminal and triple-negative cell lines in breast cancer. The metastatic potentials of HER2-positive breast cancer cell lines remain unclear.

HER2 is overexpressed in 20% of breast cancers, and HER2-positive breast cancer is known to be aggressive and have poor outcomes [4]. Although treatments targeting HER2 by chemotherapy and trastuzumab therapy has been well developed, approximately 25% of patients still experience a relapse in distant metastatic organs [5, 6]. Thus, the molecular mechanisms of metastasis in HER2-positive breast cancer must be understood, and the therapeutic strategies for metastasis should be established. However, only few *in vivo* metastasis models of HER2-positive breast cancer are available for the study of their metastatic mechanisms. The *in vivo* transplantation methods affect the evaluation of metastatic potentials and extracted metastasis gene signatures from human cancer cell lines [7]. Therefore, not only intracardiac injection, but also various transplantation methods must be used to evaluate metastatic activities.

In our previous research, we transplanted nine HER2-positive breast cancer cell lines in the brain using intracranial injection and classified them into two groups according to their proliferation abilities in the brain [8]. In this study, we evaluated the lung and bone metastatic potentials of the nine HER2-positive breast cancer cell lines by intravenous injection (IVI) and intracaudal arterial injection (CAI). As a result, the HER2-postive cell lines were classified according to their metastasis abilities. Furthermore, an expression analysis of the cell lines identified the cancer subtype and organ-specific gene signatures.

## Materials and Methods

### Cell culture

MDA-MB-453, UACC-893, HCC-2218, HCC-1419 (ATCC, Manassas, VA, USA), and ZR-75-1 cells (Institute of Development, Aging and Cancer [IDAC], Miyagi, Japan) were cultured in Roswell Park Memorial Institute medium (RPMI-1640, Fujifilm Wako Pure Chemical Corporation, Osaka, Japan) supplemented with 10% fetal bovine serum (FBS; Nichirei Biosciences Inc., Tokyo, Japan), 100 U/mL penicillin (Meiji-Seika Pharma Co., Ltd., Tokyo, Japan), and 100 μg/mL streptomycin (Meiji-Seika Pharma), and incubated under 37°C with 5% CO_2_. MDA-MB-361 and HCC-202 cells (ATCC) were cultured in RPMI-1640 supplemented with 15% heat-inactivated FBS, 100 U/mL penicillin, and 100 μg/mL streptomycin at 37°C with 5% CO_2_. BT-474 and UACC-812 (ATCC) were cultured in Dulbecco’s modified Eagle’s medium (Fujifilm Wako Pure Chemical Corporation) supplemented with 10% heat-inactivated FBS, 15% glucose, 100 U/mL penicillin (Meiji Seika Pharma), and 100 μg/mL streptomycin (Meiji-Seika Pharma) at 37°C with 5% CO_2_. UACC-812, MDA-MB-361, HCC-202 cells expressing luciferase were established by infection with lentivirus vector (pLenti-PEF1-*luc2*-IRES-BlaR). UACC-893, MDA-MB-453, HCC-2218, ZR-75-1, BT-474 and HCC-1419 cells were firstly infected with lentivirus vector (pLenti-Pubc-*mSlc7a1*-IRES-HygR) in order to express the ecotropic receptor. These 6 cell lines expressing the ecotropic receptor were infected with retrovirus vector (pMXd3-PEF1-*luc2*-IRES-BlaR). All cell lines and infection protocols were established in a previous study [8].

### Animal studies and bioluminescent imaging

Each cell line (5.0 × 10^5^ cells/100 μL phosphate-buffered saline [PBS]) was transplanted into 6-week-old female NOD.CB-17-Prkdc<scid>/J mice (NOD/scid, Charles River Japan, Inc.) via CAI or IVI [9–11]. The mice were anesthetized with 2.5% isoflurane (Fujifilm Wako) during transplantation and bioluminescence imaging (BLI). Bone and lung metastases were monitored using BLI with an IVIS Lumina XRMS In Vivo Imaging System (PerkinElmer) once a week. Each mouse was intraperitoneally injected with 3-mg D-luciferin (Gold Biotechnology Inc.) in 200-μL PBS before observation. Bioluminescent signal was measured with binning and F/stop ranges suited to each bioluminescence level. The lungs were harvested from the mice at 8 weeks after transplantation. The *ex vivo* observation of the lung was performed using IVIS-XRMS with the D-luciferin solution.

### Microarray Analysis

DNA microarray data provided in the previous research were used for genetic analysis [12]. The heatmap was drawn using the “pheatmap” package of R version 3.6.1. The Gene Ontology (GO) term enrichment analysis was performed using Metascape [13]. A principal component analysis (PCA) was performed in R with the “scatterplot3d” package. A Venn diagram was drawn using the “ggplot2” package.

### Survival analysis

Survival analysis was performed using the Kaplan-Meier method for patients with breast cancer in the Molecular Taxonomy of Breast Cancer International Consortium (METABRIC) data set, as described previously [8, 14, 15].

### Data availability

The microarray data of the nine HER2-positive cell lines were obtained from a previous study [12].

## Results

### Bone metastasis profiles of the HER2-positive breast cancer cell lines

Nine HER2-positive breast cancer cell lines expressing the *luc2* gene, namely UACC-893, MDA-MB-453, HCC-2218, BT-474, ZR-75-1, UACC-812, MDA-MB-361, HCC-202, and HCC-1419, were transplanted into NOD-SCID mice using the CAI method. After the CAI, MDA-MB-453, UACC-893, and HCC-202 proliferated rapidly at week 8 (Fig. 1a). HCC-2218 and HCC-1419 showed no tumor formation at week 8, which suggests that both have no bone metastatic potential. On the other hand, BT474, ZR-75-1, UACC812, and MDA-MB-361 migrated and survived in bone microenvironment. Their proliferation abilities were much milder than those of MDA-MB-453, UACC-893, and HCC-202 (Fig. 1b). We further classified the cell lines according to their number of metastasis tumors and proliferation ability into high, medium, low, and not applicable (N/A) groups (Table 1). We noticed that the tumor sizes of HCC-202 and MDA-MB-361 decreased after week 6. This result suggested that HCC-202 and MDA-MB-361 cells might not able to survive in long-term metastasis.

**Fig. 1.**
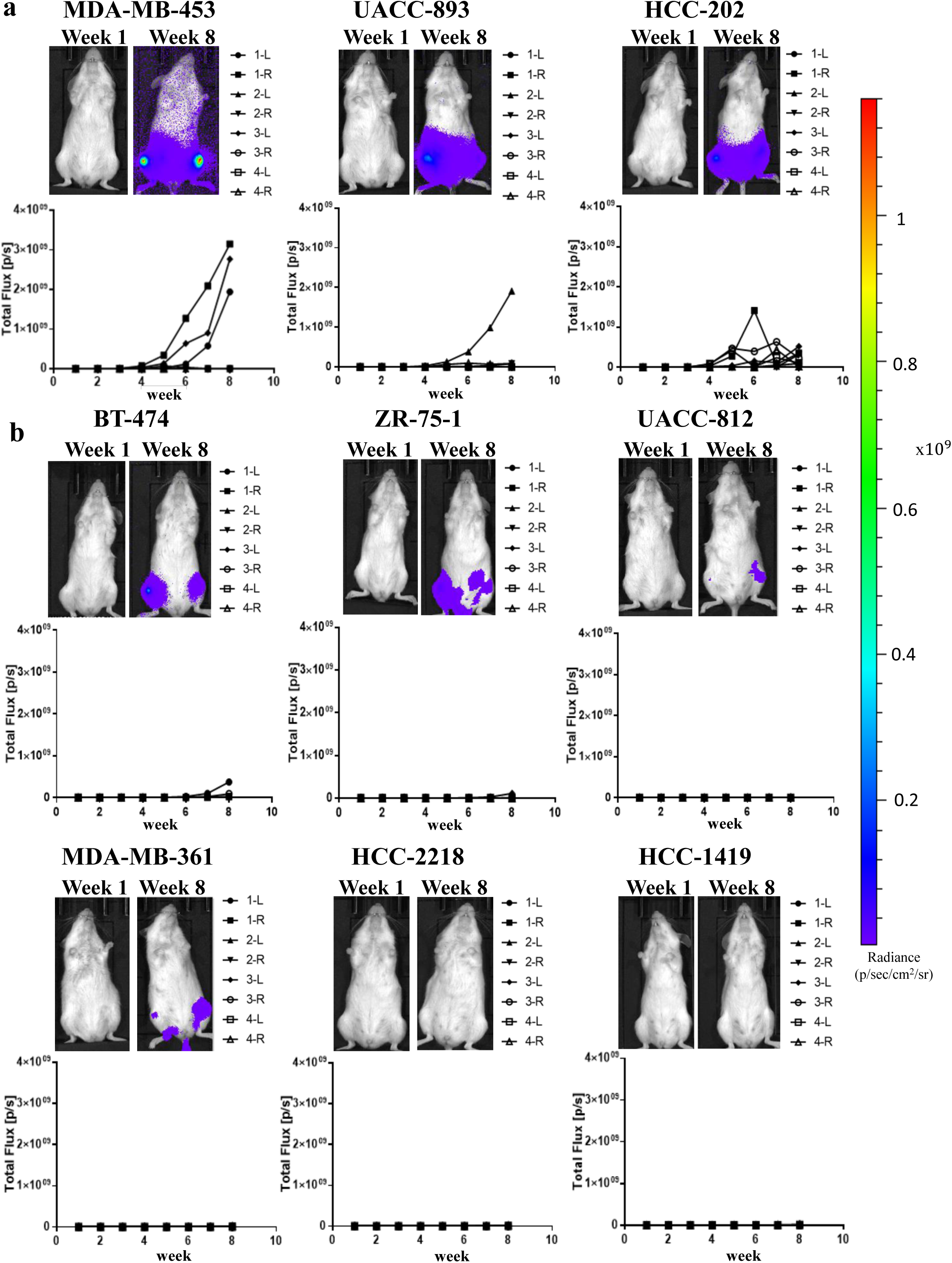
Intra-caudal arterial injection (CAI) transplantation of HER2-positive cell lines. MDA-MB-453, UACC-893, HCC-202, BT-474, ZR-75-1, UACC-812, MDA-MB-361, HCC-2218, and HCC-1419 cells were injected in NOD/SCID mouse (n = 4) by using the CAI method. The tumor growth was quantified by measuring bioluminescence every week and plotted into a growth curve. Each line showed the corresponding bone metastasis tumor. Left: Bioluminescence on week 1. Right: Bioluminescence on week 8. (a) High bone metastasis potential group. (b) Medium and low bone metastasis potential group.

**Table 1.** Metastasis profile of HER2-positive cell line

| Table 1. Metastasis profile of HER2-positive cell line |  |  |  |  |
| --- | --- | --- | --- | --- |
| Cell line | Bone metastatic tumor number / total leg number | Bone metastasis ability evaluation | Lung metastatic tumor number / total mouse number | Lung metastasis ability evaluation |
| MDA-MB-453 | 3/8 | High | 0/9 | N/A |
| UACC893 | 4/8 | High | 1/7 | Low |
| HCC202 | 7/8 | High | 2/5 | Low |
| BT474 | 3/8 | Medium | 0/4 | N/A |
| ZR-75-1 | 4/8 | Medium | 0/4 | N/A |
| MDA-MB-361 | 5/8 | Low | 0/3 | N/A |
| UACC812 | 4/8 | Low | 0/4 | N/A |
| HCC1419 | 0/8 | N/A | 0/4 | N/A |
| HCC2218 | 0/8 | N/A | 0/4 | N/A |

We divided the nine HER2-positive cell lines into two groups. The high metastatic potential group included MDA-MB-453, UACC-893 and HCC-202, and the low/no metastatic potential group included the BT-474, ZR-75-1, UACC-812, MDA-MB-361, HCC-2218, and HCC-1419 cell lines. The gene expression level was standardized into a *z*-score. The genes with average *z*-scores > 1.0 and < −1.0 were counted as upregulated and downregulated genes, respectively (Fig. 2a). We calculated the average log fold-change (FC) ratio change between the high and low potential groups. The genes with an average logFC of >1.0 or 1.0 were counted as differential expression genes (DEGs). Seventy-three upregulated genes and 69 downregulated genes were extracted as gene signatures in HER2-positive breast cancer (Fig. 2b, Table 2). The gene signatures clustered nine breast cancer cell lines into high, medium/low, and N/A metastatic potentials, consistent with our previous data (Table 1). Genes that were reported as metastasis signatures such as tumor-associated calcium signal transducer 2 (*TACSTD2*) and galectin-1 (*LGALS1*) were extracted [16, 17]. Moreover, the metabolisms of amino acids and their derivatives were mostly enriched in the high metastatic group, and the transcriptional regulation of runt-related transcription factor 3 (*RUNX3*) was overall enriched in the high potential group (Fig. 2c). Other than *RUNX3*, nuclear factor-kappa B (*NF-κB*) and metastasis-related runt-related transcription factor 1 (*RUNX1*) signals were also enriched in the high potential groups [18, 19].

**Fig. 2.**
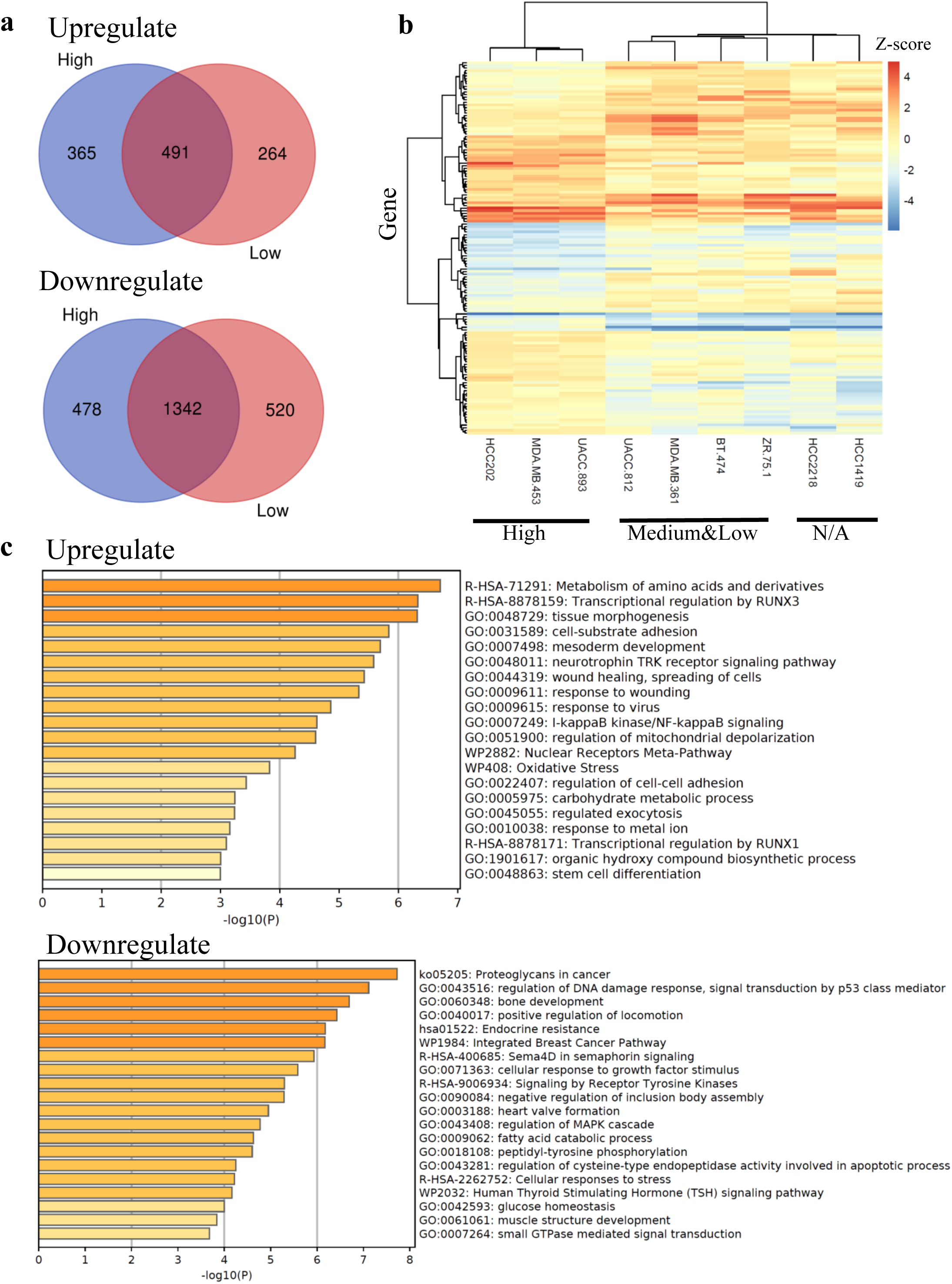
Extraction of bone metastasis gene signatures of the HER2-positive cell lines. (a) Genes were extracted and further analyzed: logFC < −1 or logFC > 1.0, p < 0.05 and FDR < 0.05. The number of upregulated and downregulated genes in the high and low potential groups are summarized in the Venn diagrams. (b) The differential expression gene (DEGs) extracted from the high and low potential groups were analyzed by hierarchical clustering and are shown in a heatmap. (c) The Gene Ontology enrichment analysis of upregulated and downregulated DEGs.

**Table 2.**
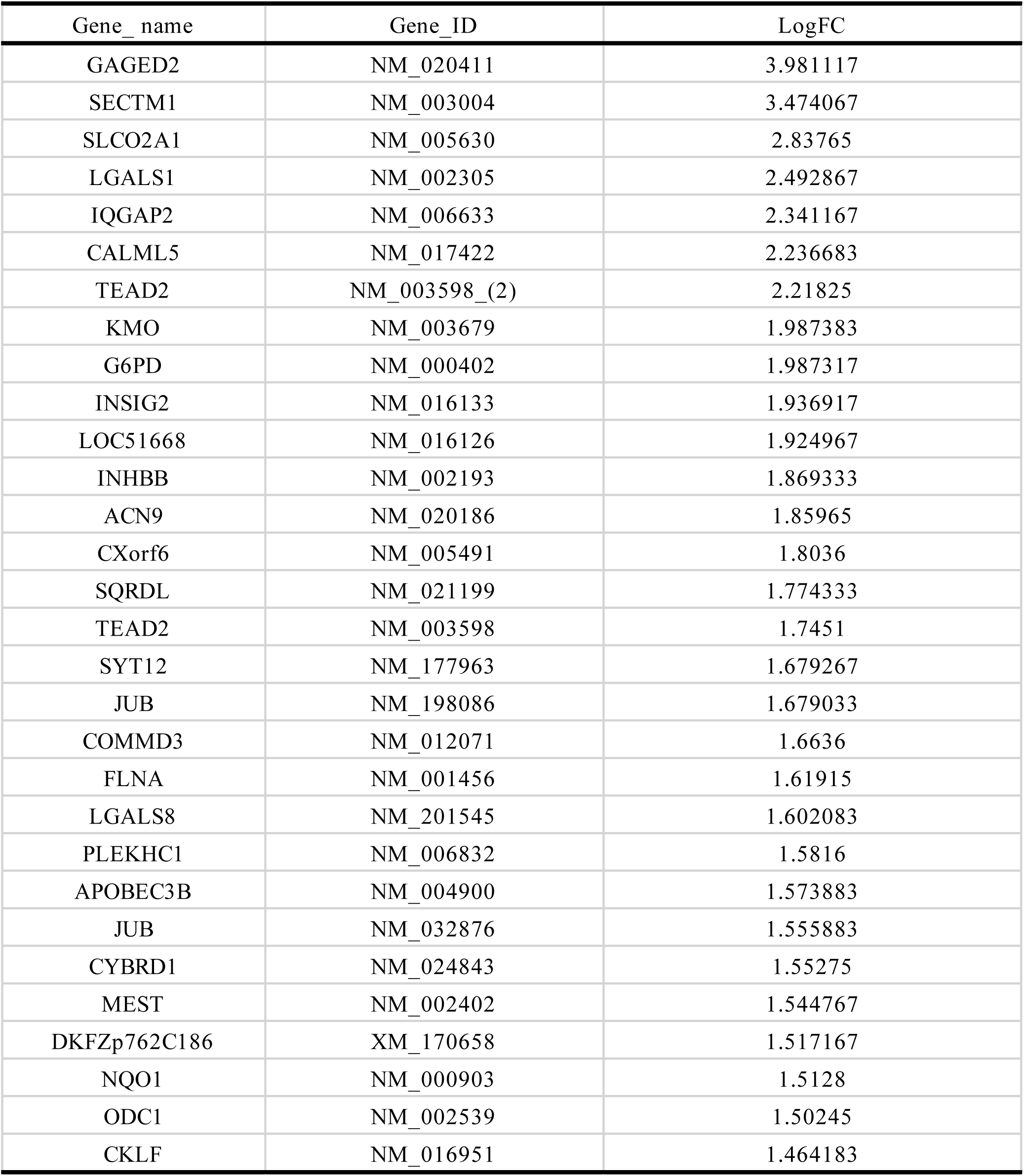
Top 30 upregulated DEGs of HER2-positive bone metastasis gene signatures

### Survival analysis of bone gene signatures

To further evaluate the relevance between the extracted gene signatures and the clinical prognosis of patients with HER2-positive breast cancer, we performed a survival analysis using the METABRIC data set (Fig. 3a, b). The upregulated gene signatures included insulin-induced gene 2 (*INSIG2*), NAD(P)H dehydrogenase, quinone 1 (*NQO1*), and the downregulated gene 4-aminobutyrate aminotransferase (*ABAT*), which correlated with the poor prognosis in HER2-positive breast cancer and all breast cancer subtypes (Supplementary Fig. S1). On the other hand, myotubularin-related protein 2 (*MTMR2*) had HER2-positive-specific clinical signatures that showed no relationship with patient prognosis in all breast cancers. This result also suggests that the clinical markers varied between HER2-positive breast cancer and other breast cancer subtypes.

**Fig. 3.**
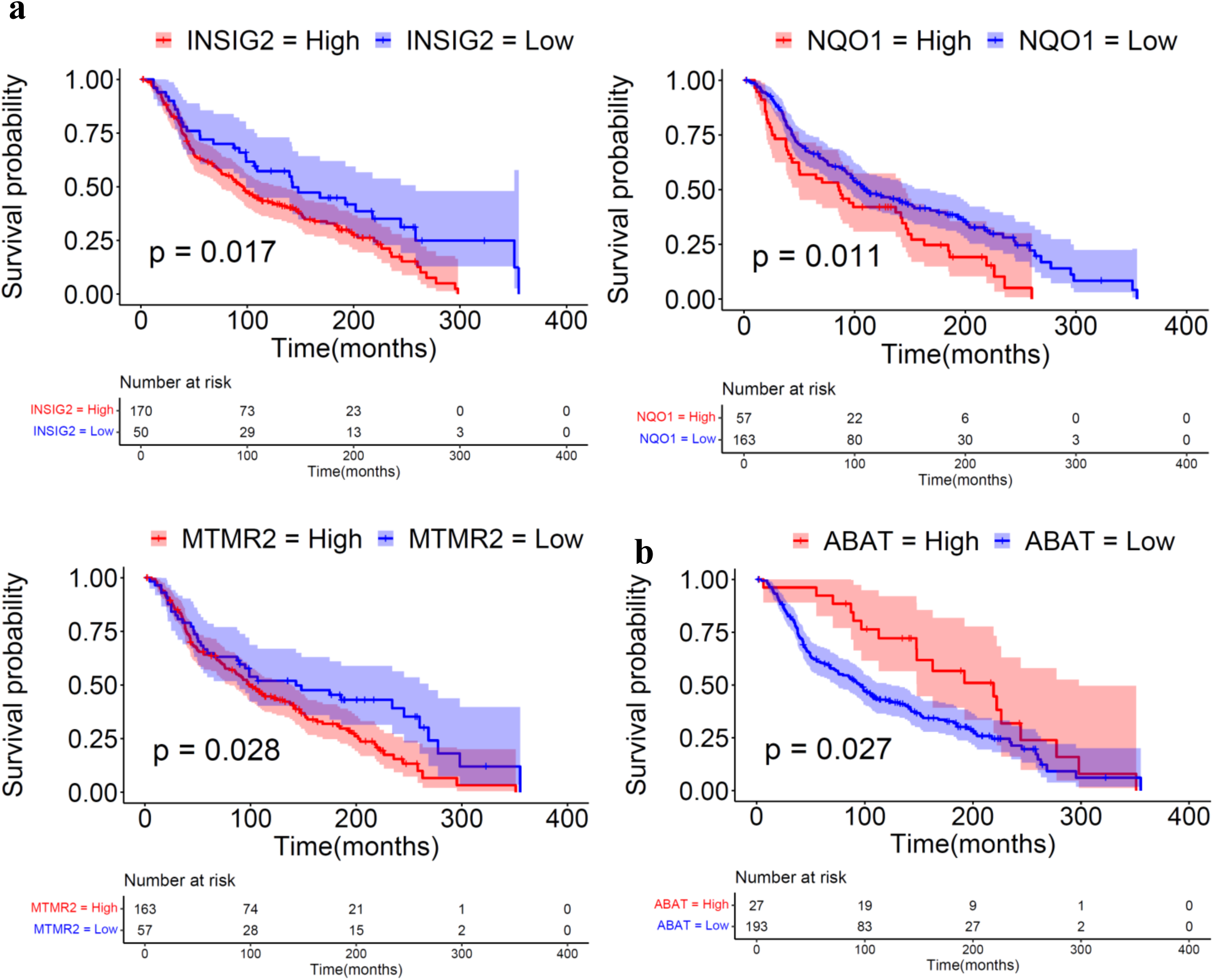
Clinical prognosis of bone metastasis gene signatures. (a) Survival analysis of differential expression gene (DEGs) in the high potential group using the METABRIC data set. The colored region along the curve shows the 95% confidence intervals. The table at the bottom lists the number of patients with high or low gene expression in the HER2-positive subtype. The results of the survival analysis of the upregulated DEGs in the HER2-positive patients are also shown. (b) Survival analysis results of the downregulated DEGs in the HER2-positive patients.

### Lung metastasis profiles of the HER2-positive breast cancer cell lines

Nine cancer cell lines were transplanted into mice via IVI. Slight but substantial luminescence was detected for two cell lines, UACC-893 and HCC-202, which suggests that they have low lung metastasis or viability in the lung (Fig. 4a). However, no luminescence was detected in the other seven cell lines, namely MDA-MB-453, ZR-75-1, HCC-1419, HCC-2218, BT-474, and MDA-MB-361, until week 8 (Fig. 4b). *Ex vivo* BLI was then performed by removing the lungs from each mouse to examine the lung metastatic ability more accurately. In the mice transplanted with UACC-893 and HCC-202, luminescence was detected from the lungs (Fig. 4c). Even *ex vivo* imaging, however, did not detect luminescence from the lungs for the other seven cell lines, namely MDA-MB-453, ZR-75-1, HCC-1419, HCC-2218, BT-474, and MDA-MB-361. On the basis of these results, we classified the UACC-893 and HCC-202 cell lines into a group with low lung metastatic potential, and the remaining seven cell lines into a group without metastasis potential (Table 1).

**Fig. 4.**
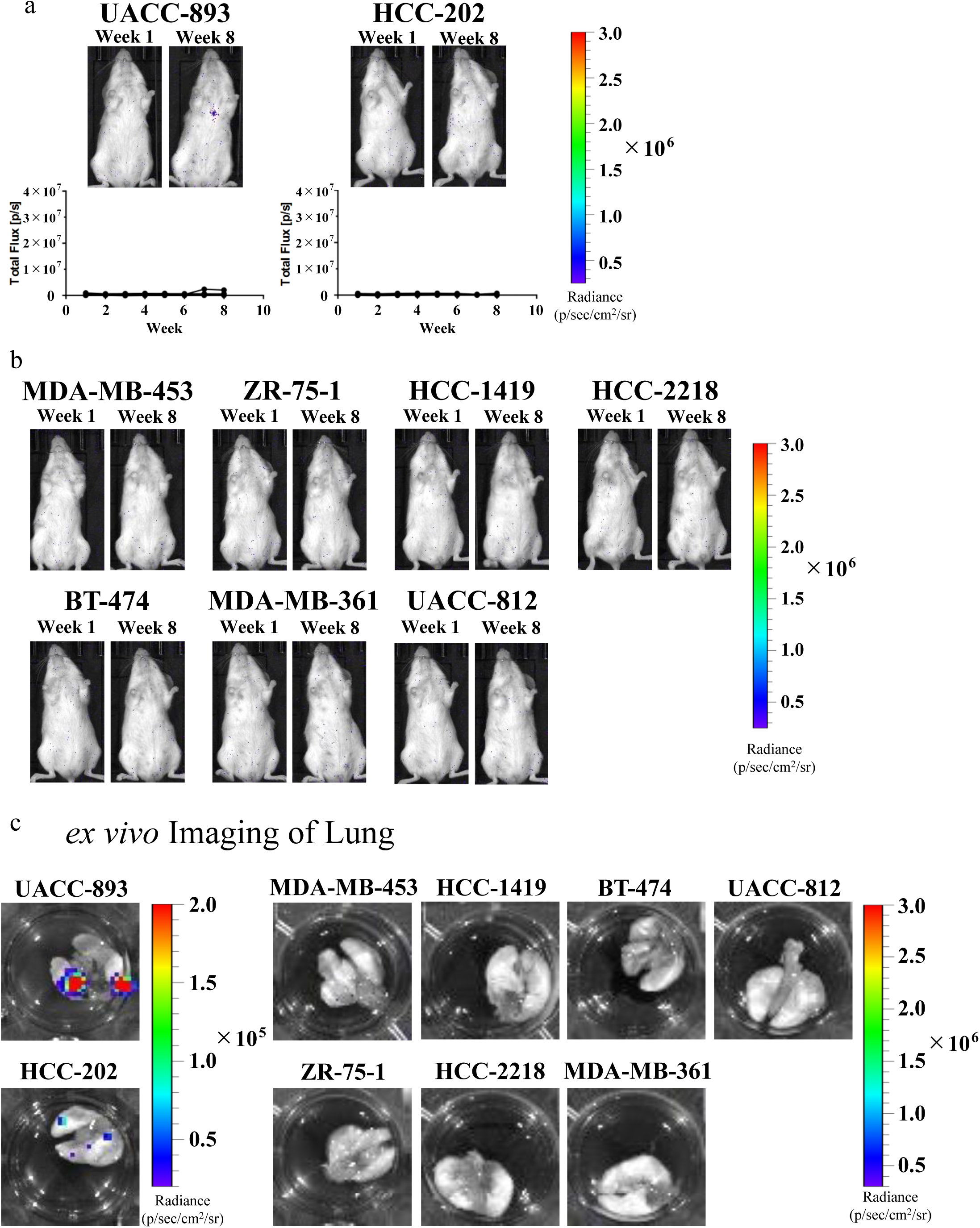
Intravenous injection (IVI) transplantation of the HER2-positive cell lines. UACC-893, HCC-202, MDA-MB-453, ZR-75-1, HCC-1419, HCC-2218, BT-474, MDA-MB-361, and UACC-812 cells were intravenously injected in the NOD-SCID mice (MDA-MB-453, n = 9; UACC-893, n = 7; HCC-202, n = 5; MDA-MB-361, n = 3; others, n = 4). Lung metastasis was quantified by measuring bioluminescence every week, and the data were plotted into a growth curve. Each cell line shows the corresponding mouse. Left: Bioluminescence on week 1. Right: Bioluminescence on week 8. (a) Cell lines in the low metastasis group. (b) Cell lines in the no metastasis group. (c) The lungs were removed from each mouse after the 8-week measurement, and luminescence from the lung was detected using *ex vivo* BLI.

From the IVI result, we reanalyzed the microarray data using the same strategy as in the reanalysis of data from the group with bone metastasis. The average *z*-score of each gene was calculated for two groups and visualized as a Venn diagram (Fig. 5a). Then, the genes highly or lowly expressed in the low metastasis group only were subjected to GO enrichment analysis. The genes related to lipid metabolism were enriched in the low metastasis group. Next, the genes with an average logFC > 1.0 or −1.0 were extracted as DEGs. In the UACC-893 and HCC-202 cell lines, 162 genes were upregulated, and 95 genes were downregulated as compared with the other seven cell lines (Fig 5b, Table 3). Among the genes that were highly expressed in UACC-893 and HCC-202 as compared with the no metastasis group are genes such as transmembrane 4 superfamily member 1 (*TM4SF1*) and *LGALS1. TM4SF1* was previously reported to promote metastatic activation in multiple organs, including the lung, across breast cancer subtypes [20]. *LGALS1* was previously reported to promote lung metastasis of claudin-low breast cancers [17].

**Fig. 5.**
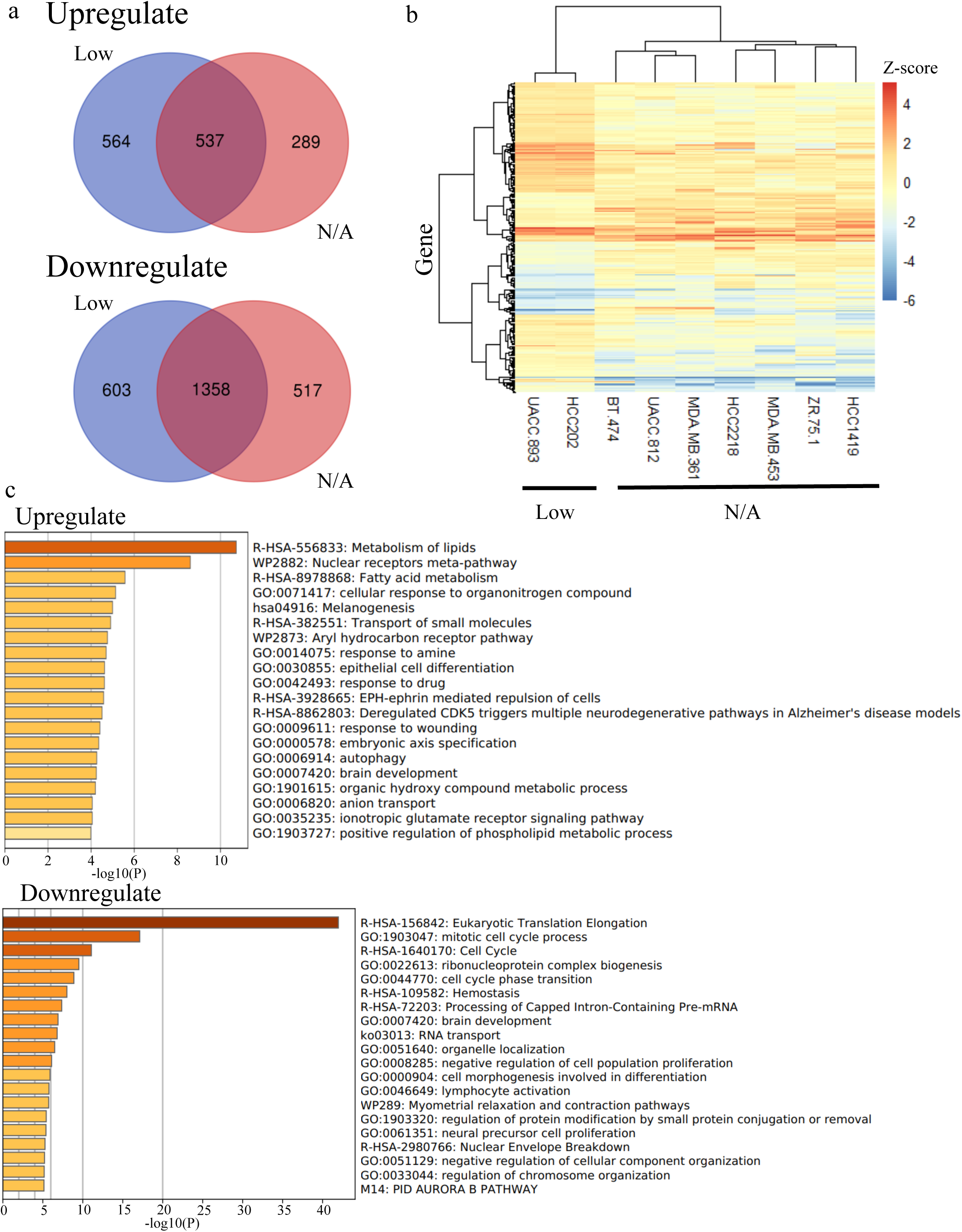
Extraction of lung metastasis gene signatures of the HER2-positive cell lines. (a) Gene signatures associated with lung metastasis or survival in the low-metastasis group were extracted in the same method as that for gene signatures associated with bone metastasis. The numbers of upregulated and downregulated genes in the low or no metastasis group are shown as Venn diagrams. (b) Differential expression gene (DEGs) between the low and no metastasis groups according to microarray data were analyzed using hierarchical clustering as a heatmap. (c) A Gene Ontology enrichment analysis of the DEGs was performed for the low metastasis group.

**Table 3.** Top 30 upregulated DEGs of HER2-positive lung metastasis gene signatures

| Gene_name | Gene_ID | LogFC |
| --- | --- | --- |
| MAGEA2 | NM_175743 | 4.4271 |
| MAGEA9 | NM_005365 | 3.6886 |
| FADS2 | NM_004265 | 3.6285 |
| MESP1 | NM_018670 | 3.4448 |
| D2S448 | XM_056455 | 3.1991 |
| CDH3 | NM_001793 | 2.934 |
| TMPRSS2 | NM_005656 | 2.8641 |
| CRIM1 | NM_016441 | 2.7695 |
| PDE4B | NM_002600 | 2.5558 |
| EDN1 | NM_001955 | 2.5277 |
| EFEMP1 | NM_004105 | 2.4872 |
| ABCC3 | NM_003786 | 2.4322 |
| PHYH | NM_006214 | 2.3426 |
| FLJ22671 | NM_024861_(2) | 2.3006 |
| LGALS1 | NM_002305 | 2.2823 |
| TEAD2 | NM_003598_(2) | 2.1974 |
| MDS025 | NM_021825 | 2.1381 |
| TP53BP2 | NM_005426_(2) | 2.1045 |
| PPARBP | NM_004774 | 2.0872 |
| KIAA0644 | AB014544 | 2.0859 |
| UBE2L6 | NM_004223 | 2.0311 |
| PCP4 | NM_006198 | 2.0165 |
| ACN9 | NM_020186 | 1.9784 |
| CAPN2 | NM_001748 | 1.9782 |
| AK5 | NM_012093 | 1.9767 |
| JUB | NM_198086 | 1.8751 |
| ASS | NM_000050 | 1.8692 |
| TENS1 | NM_022748 | 1.8676 |
| STAT5A | NM_003152 | 1.8587 |
| IBRDC2 | NM_182757 | 1.856 |

### Characterization of HER2-positive cell lines by cancer subtype-specific analysis

To further characterize these HER2-positive cell lines, we clustered the nine HER2-positive cell lines according to their whole-gene expression levels. The nine cell lines were divided into three groups (Fig. 6a). The clustering tree of the HER2-positive cell lines exhibited that the low-lung and high-bone metastasis cell lines UACC-893 and HCC-202 were relevant to each other, while the other HER2-positive cell lines were clustered independently according to their metastatic ability. Next, we quantified their metastasis potential according to their metastatic abilities to each organ site, including the brain metastatic activities and their *in vitro* proliferation ability [8], which has been previously obtained (Table S1). Three dimensional PCA (3D PCA) suggested that the UACC-893 cells had a significant difference in metastatic ability from the rest of the eight cell lines (Fig. 6b).

**Fig. 6.**
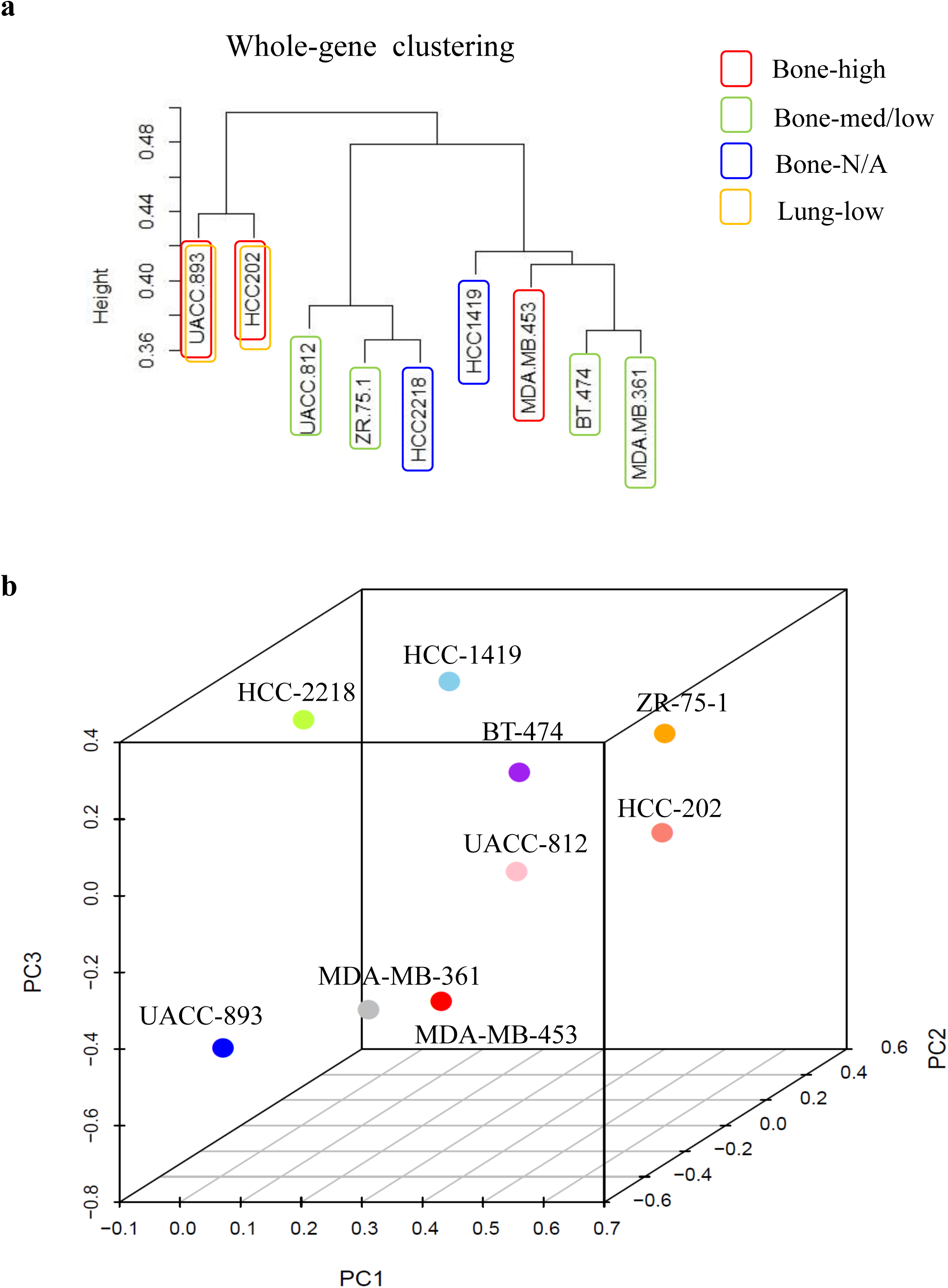
Clustering of metastatic activities in the HER2-positive cell lines. An overview of the metastatic ability of the HER2-positive cell lines is shown. (a) The clustering of whole-gene expression was performed using the wardD2 method. The cell lines were clustered into three groups. (b) The 3-D principal component analysis (PCA) plot of nine HER2-positive breast cancer cell lines according to their metastatic potentials. The PCA was performed as shown in Supplementary Table S1.

We previously established the high-metastatic cell lines of luminal breast cancer and triple-negative breast cancer (TNBC) by using the CAI method, and the bone metastasis gene signatures from luminal breast cancer and TNBC proved to be distinct from each other [10]. This suggests that the metastatic mechanism may vary among the molecular subtypes of breast cancer. Therefore, to provide a subtype-specific gene profile of breast cancer, we performed a comparative analysis of the gene signatures between HER2-positive breast cancers, luminal breast cancer, and TNBC (Fig. 7). The result showed no common upregulated or downregulated gene signature among the three subtypes. Six common upregulated signatures were found between luminal and TNBC, and three common upregulated signatures were found between luminal and HER2-positive breast cancers. On the other hand, only one common downregulated gene was found between TNBC and HER2-positive. The common downregulated signature genes both belonged to a family with disintegrin and metalloproteinase with thrombospondin motifs (ADAMTS).

**Fig. 7.**
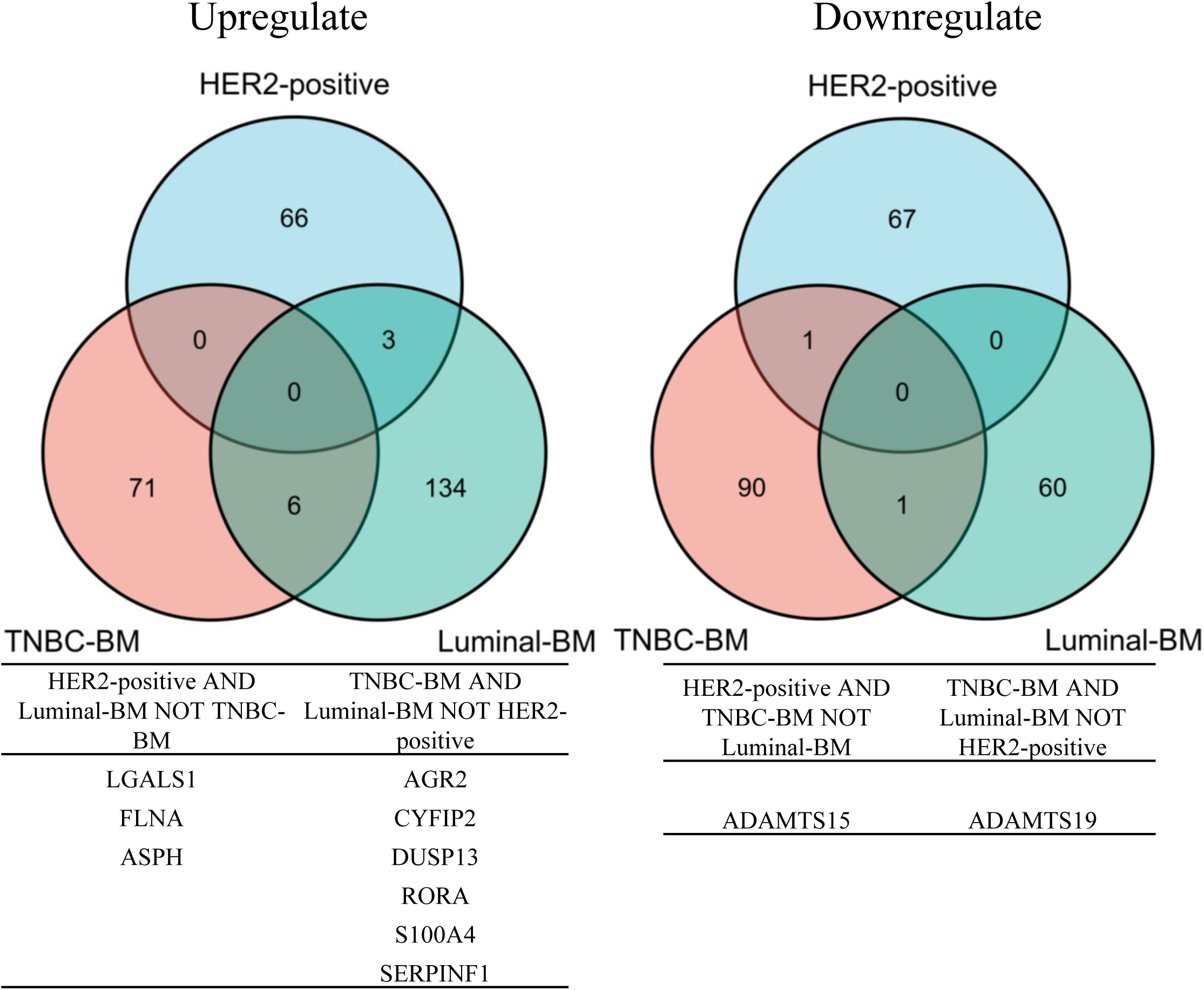
Breast cancer subtype-specific gene signatures in metastasis. The numbers of upregulated and downregulated genes among the luminal, HER2-positive, and TNBC subtypes are summarized as Venn diagrams. The commonly upregulated or downregulated genes are listed in the table at the bottom.

### Characterization of HER2-positive cell lines by organ-specific analysis

The HER2-positive cell lines exhibited various metastasis characteristics. We compared the extracted gene signatures from bone metastasis, lung metastasis, and brain colorization to demonstrate the organ-specific metastasis features. The results indicated no common upregulated signature among the three organ sites (Fig. 8). However, four common gene signatures, namely 4-aminobutyrate aminotransferase (*ABAT*), MYB proto-oncogene like 1 (*MYBL1*), Twist-related protein 1 (*TWIST1*), and olfactomedin 1 (*OLFM1*), were downregulated in the bone, lung, and brain. *ABAT* correlated with the clinical patient prognosis in both HER2-positive breast cancer and all breast cancer subtypes (Fig. 3b, Supplementary Fig S1b). The metastasis signatures of the HER2-positive cells were more commonly shared between bone and lung metastases, rather than brain metastasis. The result of the comparative analysis suggests that brain colonization of HER2-positive breast cancer had a unique mechanism compared with those in bone and lung metastases.

**Fig. 8.**
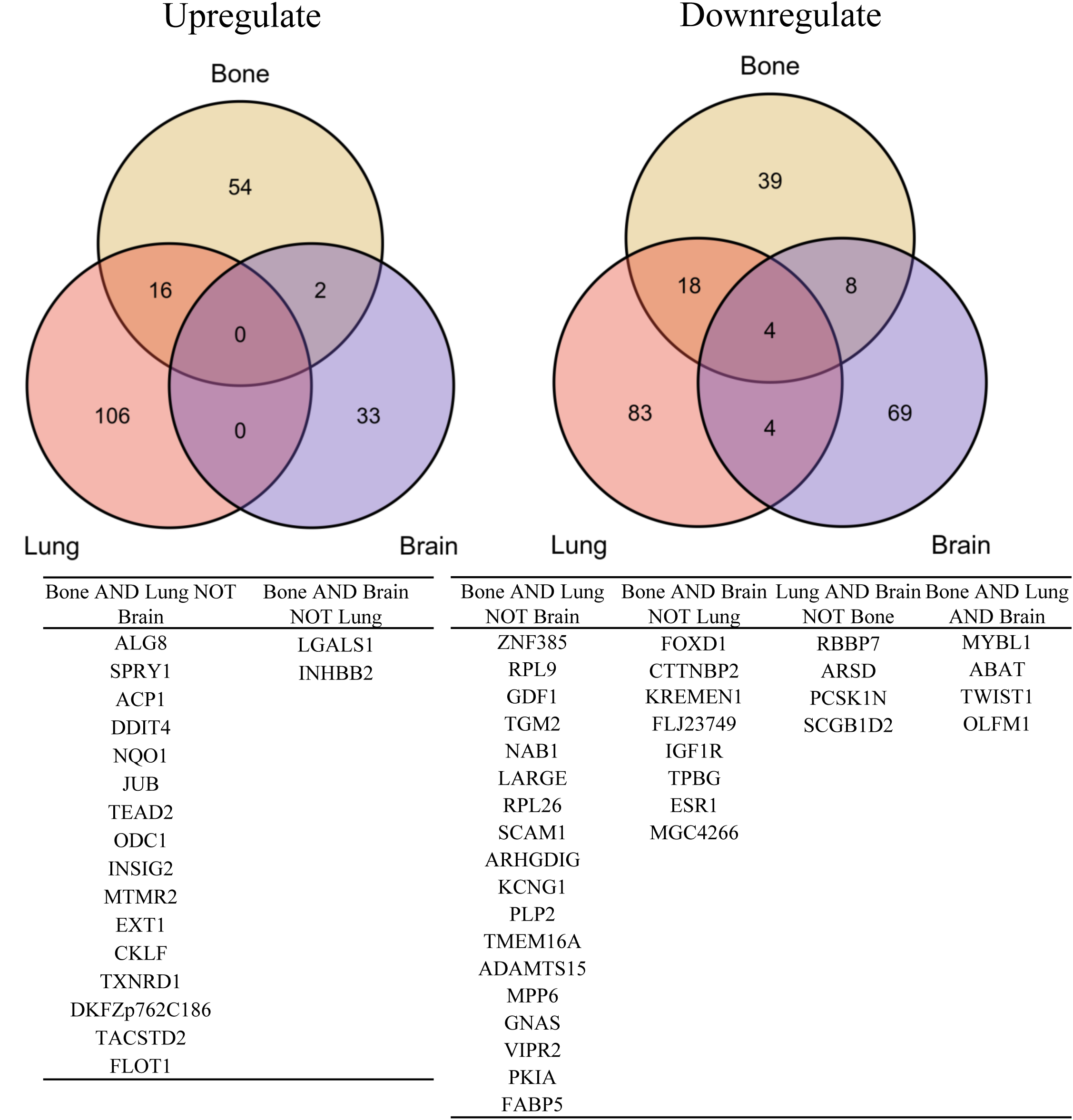
Organ-specific gene signatures in breast cancer metastasis. The numbers of upregulated and downregulated genes in the bone, lung, and brain are summarized as Venn diagrams. The commonly upregulated or downregulated genes are listed in the table at the bottom.

## Discussion

In this study, we established the xenograft models of lung and bone metastases by using nine HER2-positive breast cancer cell lines. Our results profiled their metastatic potentials in xenograft models and provided novel models for future studies on HER2-positive breast cancer metastasis mechanisms.

IVIs of UACC-893 and HCC-202 were confirmed to generate weak but significant luminescence in the lung. Thus, they can proliferate very slowly or continue to survive without proliferation. The enrichment analysis revealed that genes involved in lipid metabolism were enriched in the low metastasis group. Regulation of lipid metabolism contributes to increased aggression of breast cancer [21] and regulation of CSC [22]. In this regard, *PFKFB3* is commonly upregulated as a lipid metabolism-related gene known to be associated with cancer [23]. In addition to *PFKFB3, NDRG1* [21], *KDM5B* [24], and *JAK/STAT3* [22] have been reported as lipid metabolism-related genes associated with breast cancer. They may contribute to breast cancer malignancy through cell proliferation, migration, drug resistance, and cancer stem cells. These two cell lines may be useful for elucidating the new dormancy or survival mechanism in the lungs through the lipid metabolism. The other seven cell lines have no lung metastatic ability. Previous studies reported that IVI has low or no ability to metastasize to the lung in MDA-MB-453 and BT-474 [25]. In addition, by using intracardiac injection method, ZR-75-1 and HCC-1419 have also been reported to have no lung metastatic potential [3]. Other reports indicated that MDA-MB-453 and BT-474 cell lines were able to metastasize to the lung. However, a direct comparison is difficult because the mouse species used for transplantation were different or the number of transplanted cells was extremely large compared with that in this study [26, 27]. Moreover, although IVI was used in this experiment for the purpose of mimicking lung metastasis, previous studies confirmed that the population of enriched cells differs depending on the transplantation method [11]. In fact, one study reported that lung metastasis of the MDA-MB-453 cell line was confirmed by orthotopic transplantation in NSG mice [28]. In this study, we focused on the extravasation abilities and the following colonization and proliferation in the lung, examined using IVI. Thus, different results may be obtained if a pre-metastatic niche induced by orthotopic transplantation plays a significant role in lung metastasis.

From the results of our bone metastasis experiment, three of the nine HER2-positive cell lines, HCC-202, MDA-MB-453, and UACC-893, could rapidly grow in bone microenvironment, whereas the others could hardly form bone metastatic tumor. Even though the metastasis possibility to form a tumor in bone was different among the three cell lines, their high proliferative ability in bone microenvironment during long-term observation demonstrated that they have high bone metastatic ability. Compared with their lung metastatic ability, some of the HER2-positive cell lines exhibited stronger bone metastasis potential and brain colonization ability. Among the nine HER2-positive cell lines, UACC-893 had both lung and bone metastatic potentials and proliferation ability in the brain. This suggests that UACC-893 was a more aggressive cell line than the others and is a suitable model for multiorgan breast cancer metastasis research. As almost no study has focused on UACC-893, our result could contribute to breast cancer metastasis studies.

On the basis of the transcriptome analyses of HER2-positive cell lines, the transcriptional signals of *RUNX3* were enriched in the high bone metastasis group. *RUNX3* was a downstream effector of the transforming growth factor-β (TGF-β) signal pathway and regulates various cancer-related activities such as epithelial-to-mesenchymal transition (EMT) and cancer cell migration and invasion [29]. TGF-β is crucial in the vicious cycle within bone microenvironment and promotes the osteolytic bone metastasis of breast cancer [30, 31]. Therefore, our finding suggests that *RUNX3* signals may contribute to the vicious cycle between HER2-positive cancer cells and bone microenvironment. On the other hand, the genes that regulated the proteoglycans (PGs) of cancer, including protein kinase B (AKT1) and cell-surface glycoprotein (CD44), were downregulated in the high metastasis potential group. The PGs promote the cell-cell junction and migration ability [32] but can also suppress tumor activities [33]. They may function as tumor metastasis suppressors in HER2-positive cancer bone metastasis.

According to the subtype-specific comparison, the breast cancer subtypes (luminal, HER2, and TNBC) did not share common gene signatures. This suggests that the breast cancer subtypes have unique metastatic mechanisms. The brain colonization abilities of the HER2-positive breast cancer cell lines showed no correlation with HER2 phosphorylation or expression levels [8]. Novel factors in HER2-positive cells could possibly regulate their metastasis preference. From the organ-specific comparison analysis, four common downregulated gene signatures were extracted, including *ABAT*, whose low expression correlated with poor prognosis. *ABAT* is an inhibitory neurotransmitter in the central nervous system. Its high expression level suppressed the lung metastasis ability of MDA-MB-231 [34], and its downregulation is a hallmark of ER+ breast cancer [35]. Downregulation of *ABAT* may also contribute to the multiorgan metastasis of HER2-positive tumor cells.

In conclusion, we classified the nine HER2-positive breast cancer cell lines into metastatic subgroups according to their CAI, IVI, and transcriptomic profiles. The extracted metastasis gene signatures were potential prognostic marker genes of HER2-positive breast cancer. Our results suggest that the UACC-893 cell line is a useful model for breast cancer metastasis studies. These models and gene signatures will contribute to the further understanding of the mechanisms of metastasis in HER2-positive breast cancer.

## Acknowledgments

We thank Ms Yuka Kuroiwa and laboratory members for the meaningful comments and discussion.

## Supplementary Figure Legends

**Supplementary Fig. S1 Clinical prognosis of bone metastasis gene signatures in all the subtypes**

The survival analysis of the DEGs in the high potential group was performed using the METABRIC data set. The table at the bottom lists the number of patients with high or low gene expressions in all the breast cancer subtypes. (a) Survival analysis of the upregulated DEGs in all patients with breast cancer. (b) Survival analysis of the downregulated DEGs in all the patients with breast cancer.

